# Sex differences in the relationship between maternal and foetal glucocorticoids in a free-ranging large mammal

**DOI:** 10.1101/2023.05.04.538920

**Authors:** Bawan Amin, Ruth Fishman, Matthew Quinn, Devorah Matas, Rupert Palme, Lee Koren, Simone Ciuti

## Abstract

Maternal phenotypes can have long-term effects on offspring phenotypes. These maternal effects may begin during gestation, when maternal glucocorticoid (GC) levels may affect foetal GC levels, thereby having an organizational effect on the offspring phenotype. Recent studies have showed that maternal effects may be different between the sexes. However, how maternal GC levels relate to foetal levels is still not completely understood. Here we related, for the first time in a free-ranging large mammal, the fallow deer (*Dama dama*), maternal GC levels with foetal *in utero* GC levels. We did this in a non-invasive way by quantifying cortisol metabolites from faecal samples collected from pregnant does during late gestation, as proxy for maternal GC level. These were then related to GC levels from hair of their neonate offspring (n = 40). We have shown that maternal GC levels were positively associated with foetal GC levels, but only in female offspring. These findings highlight sex differences, which may have evolved to optimize male growth at the cost of survival.

## Introduction

Parental phenotypes can be drivers of offspring variation (Badyaev & Uller, 2009; Wolf & Wade, 2009), and in many mammalian species, these parental effects are often assumed to be mainly maternal effects because of pregnancy and nursing. Maternal effects are seen as an adaptive way in which mothers can either fine tune their offspring for the current environmental conditions (Groothuis et al., 2005; Sheriff & Love, 2013), or maximise their own reproductive output (Groothuis et al., 2019; Marshall & Uller, 2007). These effects on the offspring are present from the earliest stages of development and can have long lasting influence on offspring phenotype (Seckl & Meaney, 2004; Weinstock, 2008).

During gestation, maternal glucocorticoid (GC) levels can affect offspring phenotype in many ways, including affecting offspring birth weight and HPA-axis reactivity (Dantzer et al., 2013; Seckl & Holmes, 2007; Seckl & Meaney, 2004). These effects can linger until offspring are well into adulthood (e.g. Liu et al., 2001), which is also referred to as foetal programming (Seckl & Holmes, 2007). There is increasing evidence, however, that there are sex differences in foetal programming (Braithwaite et al., 2018; Fishman et al., 2024; Liu et al., 2001). In mice (*Mus musculus*), for instance, higher maternal GC was associated with higher foetal GCs, but only in female offspring (Wieczorek et al., 2019). In nutria (*Myocastor coypus*), *in utero* accumulated testosterone was found to be heritable, but only between parents and offspring of the same sex (Fishman et al., 2024). In birds, a meta-analysis found evidence that male offspring are typically more strongly affected by maternal steroid hormones in the egg than female offspring (Podmokła et al., 2018). Since high GC levels during gestation are associated with growth restrictions (Edwards et al., 1996; Meakin et al., 2021; Seckl & Holmes, 2007), it is possible that sex differences in foetal programming are due to the tendency of males to invest more in growth (Meakin et al., 2021).

Most studies investigating hormonal maternal effects investigate offspring GC levels after birth and most evidence comes from either clinical studies on humans (Gitau et al., 1998), rodents (Weinstock, 2008) or birds (Groothuis et al., 2019; Jenkins et al., 2014), with a paucity of data on free-ranging mammals. This is usually due to the challenges of measuring maternal and foetal hormone levels during gestation in non-captive mammals. Recent developments, however, have enabled measuring foetal hormonal levels post-parturition by quantifying steroids in neonate hair (Amin et al., 2021; Fishman et al., 2019; Kapoor et al., 2016). Hair steroid levels represent long-term levels, accumulated over weeks to months (Gormally & Romero, 2020), and therefore reflect *in utero* integrated levels when quantified in neonates (Amin et al., 2021). Whether these levels are associated with circulating maternal GC levels, however, is currently not well understood.

Here we explored, for the first time in free-ranging large mammals, the relationship between circulating maternal GC levels and offspring *in utero* accumulated GC levels. For that purpose, we collected faeces of mothers during late gestation, from which we quantified GC metabolites (Palme, 2019), and related that to their offspring hair GC levels (Amin et al., 2021), which were collected during the first days post-parturition. We ensured that females were sampled within their last five weeks of gestation, since that is when the foetus is fully covered in fur (Chapman & Chapman, 1997). To allow for sex differences in the association between maternal and foetal GC levels, we ran analyses to explore differences between male and female offspring. We had no clear *a priori* predictions regarding the existence or direction of potential sex differences.

## Methods

### Study population

We conducted this study in Phoenix Park, a 7.07 km^2^ urban park located in Dublin, Ireland. There is a resident population of free-ranging fallow deer that has been introduced in the 17^th^ century, with a population size of about 600 individuals in late summer, after fawn births (Griffin et al., 2022). Most births occur between early and late June of each year, with fallow deer does typically producing one fawn per year. Fallow deer is a hider species and fawns remain hidden, usually in tall grass or understory vegetation, away from the main doe herd during the first weeks of life after which they are brought into the doe herd by their mothers (Chapman & Chapman, 1997; Ciuti et al., 2006). Fawns are occasionally predated upon by red foxes (*Vulpes vulpes*), the only natural predator in the park, and domestic dogs who are brought into the park by public visitors.

### Faeces collection and analysis of maternal faecal cortisol metabolites (FCMs)

Faecal samples were collected between 8 AM and 3 PM, from May 19^th^ until May 29^th^ 2020, when the does were in late gestation. We made sure to sample during this period in order to compare maternal and foetal GC levels, since that is when the foetus is covered in hair (Chapman & Chapman, 1997). Groups of deer were observed from a distance of 50 meters using a spotting scope to identify individuals (>80% of the population is identifiable via unique colour coded ear-tags). Fresh faecal samples were collected within a minute of defecation and immediately stored in zip locked bags. These were kept in a cooler bag until they could be stored in a freezer at −20**°**C, which was always within a few hours. Samples (see *Sample sizes*-section below) were kept frozen on dry ice during transportation to the University of Veterinary Medicine (Vienna, Austria), where cortisol metabolites in the samples were quantified (Palme, 2019). We added 5 mL of methanol (80%) to weighed aliquots (0.5 g) of homogenized faecal samples, after which the samples were shaken and centrifuged as previously described (Palme et al., 2013). FCMs were analysed with an 11-oxoaetiocholanolone enzyme immunoassay (for details see Möstl et al., 2002), which was previously validated for fallow deer by administering an adrenocorticotropic hormone to fallow deer individuals (Konjević et al., 2011). In their study, Konjević et al. found a clear increase of FCM levels after 22 hours, indicating that this method is suitable for monitoring adrenocortical activity in our species.

### Neonate hair collection and hair GC level quantification

Neonate hair samples were collected during the fawn captures in June 2020. Fawns were routinely captured with fishing nets, typically during their first week of life, as part of the long-term management of the population. During handling, we collected physiological and behavioural data (see Amin et al., 2021 for full details on the capture protocol), as well as hair samples (>100 mg) from the belly of the fawn using an electric trimmer (Wahl model 9639; Wahl Clipper Corporation). We extracted cortisol levels from fawn hair using a standardized protocol for hair-testing (Fishman et al., 2019; Koren & Geffen, 2009), by using commercial enzyme immunoassay (EIA; Salimetrics; Ann Arbor, MI, USA, item no. 1-3002) kits. Full details on these extractions and validations are described in Amin et al. (2021) and in the supplementary material (S1).

### Mother-fawn pairing

Mother-fawn pairs were based on field observations with data collection starting in July, when young fawns were making their first entrances into the female herds. For details regarding the pairing, see Griffin et al. (2022) and Supplementary S1.

### Sample sizes

We collected and analysed a total of 164 faecal samples of 99 different pregnant does. We removed one outlier from an individual that had multiple samples, because it was far outside the range of the other values (value: 1319 ng/g faeces; range of other values: 33-751 ng/g faeces). Removing this single sample did not affect our results. For individuals with multiple values, we took the mean of these values to create a dataset that had one estimate per individual. Out of these, we were able to pair 41 does with their fawns through the mother-fawn observations during summer. We also removed one outlier from the male neonate GC dataset because it was far outside the range of the other values (value: 29.5 pg/mg hair; range of other values: 6.5-18.3 pg/mg hair). Our final sample size thus consisted of 40 fawns (18 females and 22 males) paired with their mothers. Running our analysis with and without this outlier revealed that removing this outlier did not affect the estimates of our models, although it did affect statistical significance of one of our tests due to an increased variance (see Results).

### Statistical analysis

All analyses were performed in RStudio (Version 1.3.1093) using R version 4.0.2 (R Core Team, 2020). For the maternal FCM levels, we checked among-individual repeatability with the *rptR*-package (version 0.9.22; Stoffel et al., 2017). We then first checked whether maternal FCM or neonatal GC levels differed between the offspring sexes by performing a t-test (that considered unequal variances). Next we explored, for each sex separately, the relationship between neonate hair GCs and maternal FCMs through the use of linear models (see details below). We were at first reluctant to run one model with both sexes including an interaction term. This was because models with interaction terms require high sample sizes (Harrison et al., 2018). After analysing the separate models, however, we considered it useful to further investigate whether the slope between neonatal GCs and maternal FCMs were different for the two foetal sexes *post hoc*. We therefore ran the additional model, including both sexes (see details below).

To investigate the relationship between neonate hair GCs and maternal FCMs, we ran a linear model for each fawn sex separately. In both models, we had the neonate hair GCs as the response variable and the maternal FCMs as the explanatory variable. Maternal FCM levels were log-transformed to improve model fit, since these suffered from a slight positive skew. Model assumptions were checked with the *DHARMa-*package (Version 0.4.3; Hartig, 2021) and were successfully met (see R-script from https://doi.org/10.5281/zenodo.8355167). Statistical inferences were made based on the estimate and the associated 95% Confidence Interval from the models. During the preliminary analysis, we included the number of days between the collection of mothers’ faeces and the day of fawn birth as an explanatory variable in our models. This was to account for potential variation in FCM levels as a function of gestational day of collection (i.e. mid-May vs late May when closer to parturition). This additional predictor had no clear effect and made our models worse, indicated by a higher AIC in all cases (ΔAIC_females_ = 1.99, ΔAIC_males_ = 0.54, ΔAIC_both_ = 1.43); we therefore decided not to include it. We plotted our results using the *ggplot2*-package (Wickham, 2016).

Finally, we decided to run an additional model *post-hoc*, including both sexes in the same model, to further investigate whether the slopes of our models were different between male and female offspring. We ran a linear model with neonate hair GCs as the response variable. As explanatory variables, we included maternal FCMs (log transformed), fawn sex and the interaction between both these explanatory variables.

## Results

Considerable variation was found in both the individual maternal FCM levels (range: 91-608 ng/g faeces) as well as the neonate hair GC levels (range: 6.50-18.30 pg/mg hair; outlier value: 29.5 pg/mg hair). Repeatability of FCM levels was low (R = 0.126, 95% CI [0, 0.351], p = 0.07), indicating that there was quite some within-individual variation. We found no clear sex differences, neither for mean maternal FCM levels (t-test; t = 0.79, 95% CI [-40.68, 93.11], p = 0.43) nor for mean neonate hair GC levels (t-test; t = 0.26, 95% CI [-1.74, 2.25], p = 0.80).

We found that female hair cortisol was positively associated with maternal FCMs (LM: β = 5.88, 95% CI [0.20, 11.57], p = 0.04, N = 18, R^2^_adjusted_ = 0.18; Fig. 1A). Male hair cortisol levels were not associated with maternal FCMs (LM: β = 0.67, 95% CI [-1.30, 2.64], p = 0.49, N = 22, R^2^_adjusted_ = −0.02; Fig. 1B). Our *post-hoc* model confirmed that there was indeed an interaction between foetal sex and maternal levels, where the slope between maternal and foetal levels was lower in male offspring compared to female offspring (LM: β = −5.21, 95% CI [-10.34, −0.08], p = 0.047, N = 40, R^2^_adjusted_ = 0.10). Including the outlier that was removed did not affect the estimate. However, it did increase the variance of the interaction and thus, the interaction was no longer statistically clear (LM: β = - 5.41, 95% CI [-12.49, 1.68], p = 0.13, N = 40, R^2^_adjusted_ = 0.02).

**Figure 1:**
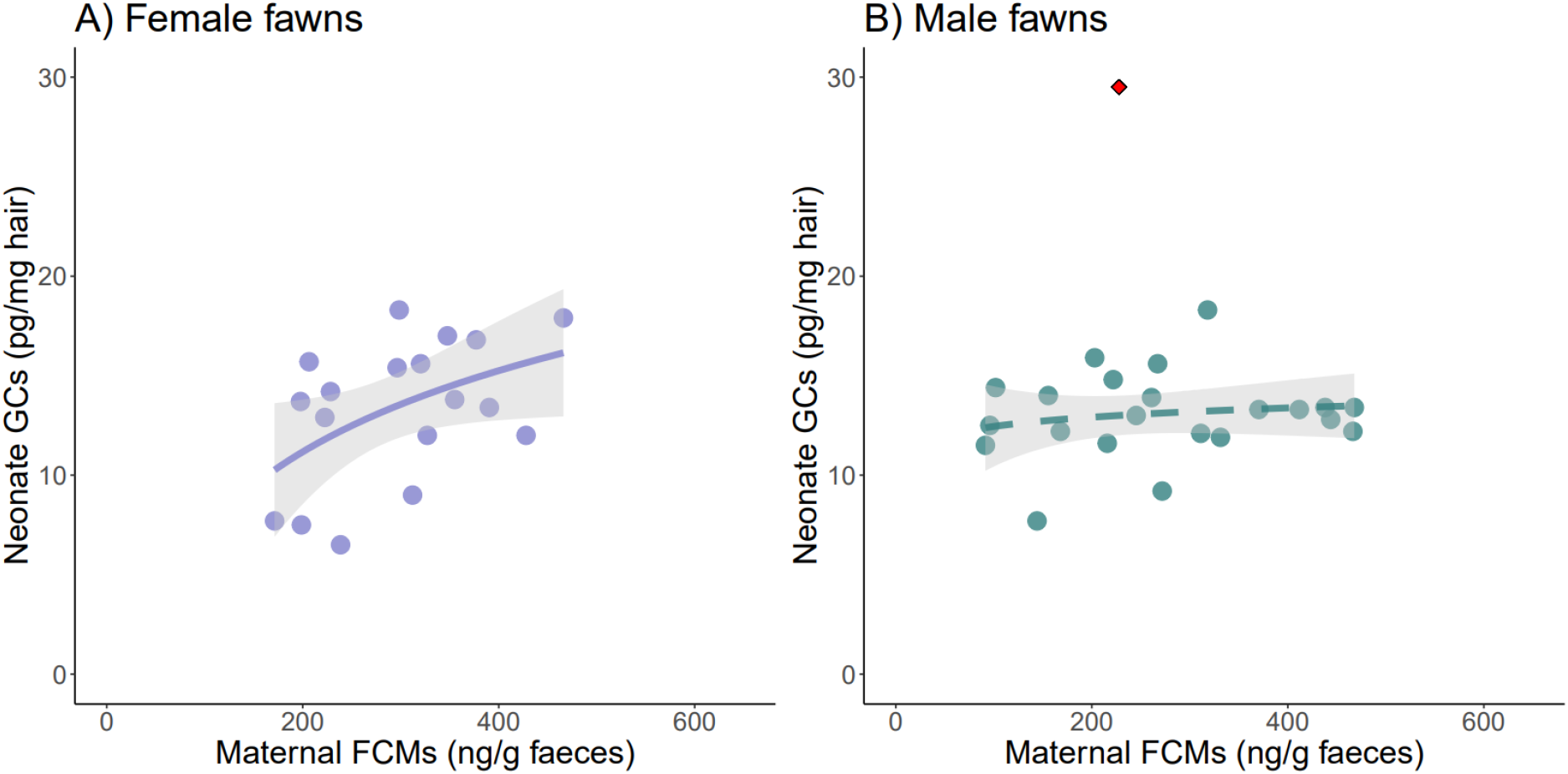
The relationship between maternal faecal cortisol metabolites (FCMs) and neonate hair GCs, for female (A) and male (B) offspring. Trendlines and their 95% confidence intervals are fitted in each plot. Solid trendline indicates a statistically clear relationship. The neonate hair GC outlier that was omitted from our analysis is indicated with a red diamond in panel B.

## Discussion

In this study we found that maternal FCM levels were positively related to foetal GC levels, but only in females. In male offspring, we found no clear relationship between maternal and foetal GC levels. These findings suggest that maternal GC levels may affect male offspring less than female offspring. We discuss here how, from an evolutionary point of view, sex-specific mechanisms may have evolved to optimize life-history trade-offs.

Like many other species, fallow deer have a skewed mating distribution (McElligott et al., 2001; Moore et al., 1995). While females tend to produce one fawn per year (Chapman & Chapman, 1997), male reproduction skew is steep. Only a small proportion of the males reproduces, with the majority having no or very few offspring (Ciuti et al., 2011; McElligott et al., 2001; Moore et al., 1995). Low quality sons assumably end up costing resources, with no fitness benefits in the long run, possibly driving selection to prioritize male growth over survival. A recent study has shown that male birthweights in our populations are indeed higher than female birthweights (Griffin et al., 2023).

GC levels play a key role during gestation and have major effects on offspring phenotype early in life. In addition to being crucial for organ development during the late stages of gestation (Kitterman et al., 1981; Liggins, 1994), GCs play a role in fighting inflammation (Auphan et al., 1995) and thereby help to keep the foetus vital. At the same time, high GC levels may restrict growth (Edwards et al., 1996; Meakin et al., 2021; Seckl & Holmes, 2007). As males tend to prioritize growth (Meakin et al., 2021), there should be sex differences in the mechanisms that evolved. For example, maternal GC contribution may be limited in male foetuses to prevent growth retardation, at the cost of prenatal survival. This may contribute in some degree to higher prenatal mortality rates in males (Desportes et al., 1994; Eriksson et al., 2010; Kruuk et al., 1999).

The mechanisms underlying these sex differences may originate in the mammalian placenta. Previous studies, in rodents and humans, have shown sex differences in the structure and activity of the placenta (Meakin et al., 2021; Murphy et al., 2003; O’Connell et al., 2013; Rosenfeld, 2015; Wieczorek et al., 2019). There is also evidence that maternal GCs may transport more easily through female placentas than males’ (Wieczorek et al., 2019). This may be, for instance, through different levels of expression and activity of enzymes or transporters that mitigate different maternal GC level spill over between males and females.

One of the key enzymes active in the placenta, 11β-hydroxysteroid dehydrogenase type 2 (11β-HSD2), inactivates cortisol to cortisone (Edwards et al., 1996; Tomlinson & Stewart, 2001). Placental 11β-HSD has been frequently shown to be positively related to birthweight (Edwards et al., 1996; Meakin et al., 2021; Seckl et al., 2000; Seckl & Meaney, 2004), and likely plays a role in preventing growth restriction. In humans and mice, studies have shown that male foetuses have higher expression or activity of 11β-HSD2 in response to maternal GC surges (Murphy et al., 2003; Wieczorek et al., 2019). Similarly in sheep, male foetuses were shown to increase 11β-HSD2 in response to maternal dexamethasone administration, whereas female foetuses did not (Braun et al., 2009). This may explain why a different study found that, following an antenatal GC treatment, growth restriction was lower in males than in females (Miller et al., 2012). In addition, male mice were shown to have increased ATP-binding cassette transporters, which mediate GC efflux toward maternal circulation (Wieczorek et al., 2019), indicating that there are different mechanisms through which males may be able to limit maternal GC levels (Montano et al., 1993). This altogether may explain why females may have higher foetal survival (Meakin et al., 2021), whereas males may be larger and faster growing.

We acknowledge that there are some shortcomings to this study. We report patterns taken over only one year, whereas ecological patterns may vary between years. Furthermore, a modest sample size (due to the challenges of the design), restricted more elaborate analyses. Repeatability of the maternal FCM levels was also low, indicating ample within-individual variation. Ideally, a future study will include multiple samples of each individual to account for this within-individual variation. Nevertheless, as one of the first studies quantifying the relationship between maternal and foetal GC levels non-invasively, this study provides novel insights into how these fundamental relationships may function in a free-ranging large mammal population. It is important to note that the nature of the study described here is exploratory, meaning that it generates hypotheses which should be tested in the future.

To conclude, we explored in this study the relationship between maternal and foetal GC levels in a free-ranging population of fallow deer. Our findings suggest that there are sex differences in the underlying evolutionary processes, which may have optimized growth at the cost of survival for male offspring, whereas maternal and female offspring GCs are associated. Our study furthermore suggests, in line with previous research, that the amount of GCs that the foetus is exposed to is not fully controlled by the mother (Gitau et al., 1998), but is also influenced by the offspring (Groothuis et al., 2019). Most of the existing literature uses the rodent or human model, with very little existing studies on other taxa. Although some features of the placenta may be conserved, there are evolutionary differences between species (Fowden, 2003; Rosenfeld, 2015). Non-model species may provide different outcomes (Fishman et al., 2019), through which we can gain fundamental insights of evolutionary mechanisms. This study provides novel insights into possible sex-specific maternal effects, and a method that can be viable for other systems, enabling the study of patterns rarely studied in the wild.

## Acknowledgements

We thank the Office of Public Works (OPW), Ireland, for funding (grant no. R18625) and support. We thank the SBES in University College Dublin (UCD), and the British Deer Society (R21845) for funding this project. We also would like to thank the Higher Education Authority of Ireland for providing a Covid-19 related Research Cost Extensions to BA. The funders had no role in study design, data collection and analysis, decision to publish, or preparation of the manuscript. We declare that none of the authors have a conflict of interest.

## Open Science and data availability statement

Following Kane & Amin (2023), all versions, updates and additional material are stored transparently on OSF (DOI 10.17605/OSF.IO/4YMC8). In addition, we have published the data and the script used for the main analysis, along with the raw faecal data under CC-BY 4.0 license on Zenodo (Amin et al., 2023).

## Author contributions

BA: Conceptualization, data curation, formal analysis, investigation, visualization, writing -original draft preparation; RF: Conceptualization, writing – original draft preparation, writing – review & editing; MQ: Data curation, investigation, writing – review & editing; DM: Investigation, methodology, writing – review & editing; RP: Investigation, methodology, writing – review & editing; LK: Investigation, methodology, writing – review & editing; SC: Conceptualization, funding acquisition, resources, project administration, supervision, writing – review & editing

