## Supplementary S1 for "Sex differences in the relationship between maternal and foetal glucocorticoids in a free-ranging large mammal"

**Description of document**

In this document, we provide additional details regarding the extraction of cortisol from fawn hair and the mother-fawn pairing reported in the main text. These are discussed below.

**Fawn hair cortisol extraction**

Hair was washed twice with isopropanol and left to dry overnight. Then, it was weighed and placed in a glass vial. Methanol was added and the vials were sonicated for 30 min and then incubated overnight at 50°C with gentle shaking. The methanol was collected and evaporated under a stream of nitrogen. Samples were reconstituted in 10% methanol and 90% assay diluent that was provided with the commercial enzyme-linked immunosorbent assays (EIA; Salimetrics; Ann Arbor, MI, USA, item no. 1-3002) according to manufacturer's recommendations.

Serial dilutions of hair pool (N = 13 samples) for cortisol validation showed linearity between 5 - 60 mg hair (equivalent to 0.111 - 3 µg/dL cortisol). Dilutions of the pool were parallel to the kit standards (univariate analysis of variance in SPSS; P = 0.249). According to the manufacturer, antibody cross-reactivity was reported as 19.2 % for dexamethasone, and less than 0.6 % for all other steroids. Intra-assay repeatability using 2 duplicates of the pool (n = 4) on the same ELISA plate was 1.96 %. Inter assay precision using duplicates of the pool on 4 different plates was 17.4 %. Recovery was calculated as 117.18 % by spiking a known amount of exogenous cortisol.

**Mother-fawn pairing**

Trained observers made behavioural observations from July until December, with the bulk of the observations happening between July and September. We recorded three different types of interactions between does and fawns: 1) suckling by the fawn, where we clearly distinguished between allosuckling (from the back; not counted) and frontal suckling (where the doe can see the fawn), 2) social grooming between the doe and the fawn, 3) following behaviour, where the fawn sticks closely to the doe. To limit incorrect pairing, we only confirmed a mother-fawn pair if the same pair had at least 2 independently observed interactions. We recorded a total of 473 interactions of 88 different fawns. That led to a total number of 61 mother-fawn pairs, of which both the mother and the fawn were tagged. Two fawns were excluded from the analyses since they had at least two independent observations with two separate does.
